# Integrating natural history-derived phenomics with comparative genomics to study the genetic architecture of convergent evolution

**DOI:** 10.1101/574756

**Authors:** Sangeet Lamichhaney, Daren C. Card, Phil Grayson, João F.R. Tonini, Gustavo A. Bravo, Kathrin Näpflin, Flavia Termignoni-Garcia, Christopher Torres, Frank Burbrink, Julia A. Clarke, Timothy B. Sackton, Scott V. Edwards

## Abstract

Evolutionary convergence has been long considered primary evidence of adaptation driven by natural selection and provides opportunities to explore evolutionary repeatability and predictability. In recent years, there has been increased interest in exploring the genetic mechanisms underlying convergent evolution, in part due to the advent of genomic techniques. However, the current ‘genomics gold rush’ in studies of convergence has overshadowed the reality that most trait classifications are quite broadly defined, resulting in incomplete or potentially biased interpretations of results. Genomic studies of convergence would be greatly improved by integrating deep ‘vertical’, natural history knowledge with ‘horizontal’ knowledge focusing on the breadth of taxonomic diversity. Natural history collections have and continue to be best positioned for increasing our comprehensive understanding of phenotypic diversity, with modern practices of digitization and databasing of morphological traits providing exciting improvements in our ability to evaluate the degree of morphological convergence. Combining more detailed phenotypic data with the well-established field of genomics will enable scientists to make progress on an important goal in biology: to understand the degree to which genetic or molecular convergence is associated with phenotypic convergence. Although the fields of comparative biology or comparative genomics alone can separately reveal important insights into convergent evolution, here we suggest that the synergistic and complementary roles of natural history collection-derived phenomic data and comparative genomics methods can be particularly powerful in together elucidating the genomic basis of convergent evolution among higher taxa.

## Introduction

Convergent evolution is the independent acquisition of similar features in distantly related lineages [1]. Ever since Darwin suggested that similar traits could arise independently in different organisms [2], understanding the underlying causes and mechanisms of convergence has been one of the fundamental objectives of evolutionary biology. Convergent evolution has been a central component to study evolutionary predictability [3] by integrating phenotypic, phylogenetic, and environmental data [1,4,5]. While convergent evolution is usually presumed to be the result of adaptation [6–8], it is clear that convergent patterns can also result from non-adaptive processes such as exaptation, evolutionary constraints, demographic history [1,5,9,10] or hemiplasy [11]. Conceptual differences in defining phenotypic convergence as process-based (when trait similarity evolves by similar forces of natural selection) versus pattern-based (when lineages independently evolve patterns of similar traits, regardless of mechanism) have practical implications for the adequate identification and measurement of convergent traits [12]. In addition to such challenges for defining convergence at the phenotypic level, additional uncertainties exist for defining the genetic basis of phenotypic convergence [13,14]

Convergent phenotypes may or may not share a genetic basis at many different hierarchical levels (e.g., nucleotide, gene, protein, regulatory networks, function; Fig. 1) [5]. Additionally, high-levels of pleiotropy, already recognized as a likely component of many cases of convergence [citation], means that our definition of “genomic basis” of convergence may require expansion to include the role of individual genes participating in multiple networks as well as functionally overlapping networks that may not share many genes.

**Fig. 1.**
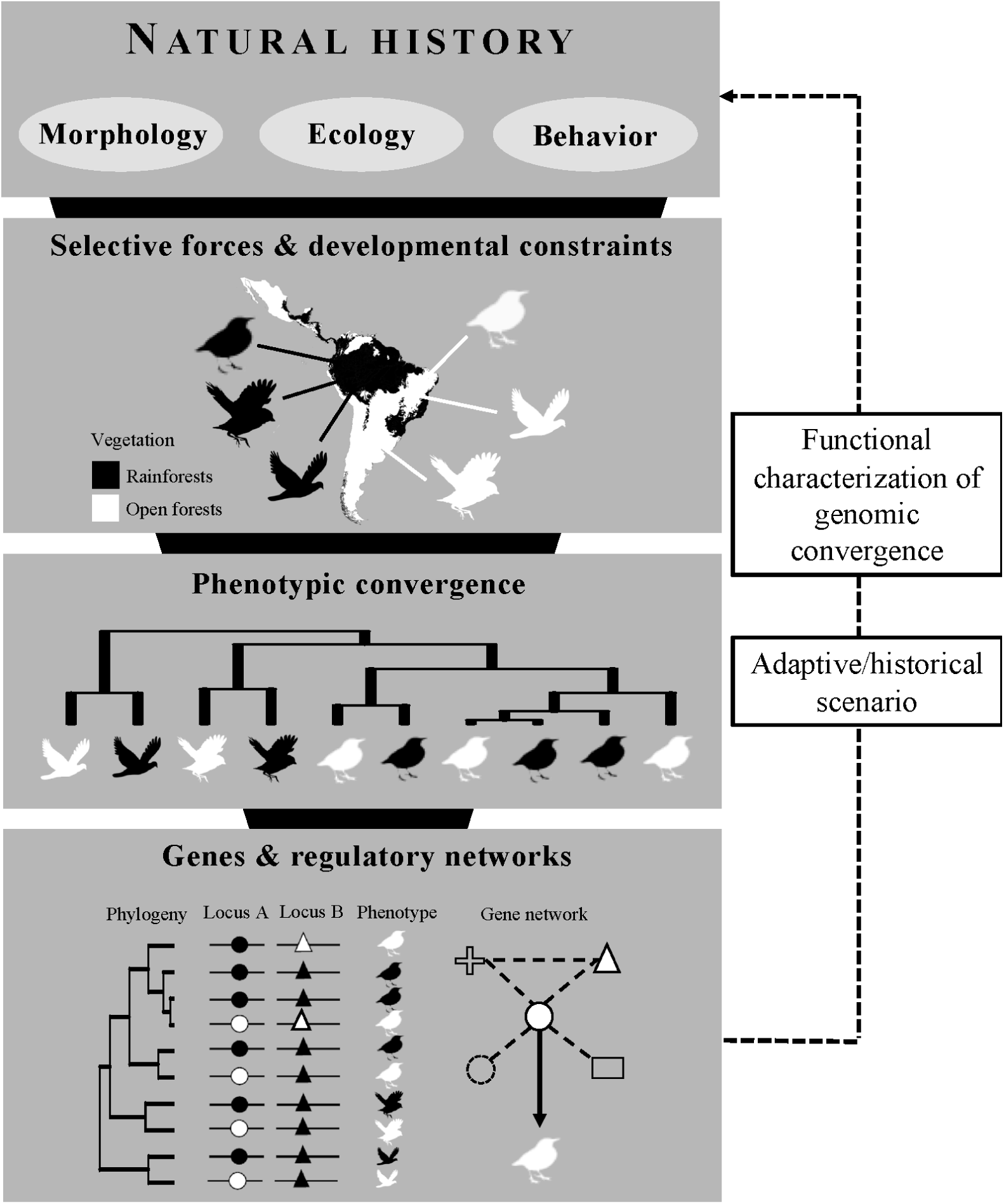
Conceptual framework for studies on the genomics of phenotypic convergence. Starting from the top, organismal expertise and knowledge of natural history are the starting points for such studies. Environmental gradients and constraints in physiology, biochemical and developmental pathways may limit or direct trait evolution, potentially driving phenotypic convergence. Phylogenetic comparative methods can be used to test, quantify and visualize instances of convergence. Finally, comparative genomics methods can be used to test whether convergent phenotypes have common underlying genomic mechanisms at various hierarchical level of individual genetic loci or regulatory networks. Functional validations of genes or pathways identified from genome-wide scans provide means to test the role of specific genomic regions in producing a given convergent phenotype and to attempt historical reconstruction of evolutionary events.

Here, we explore how recent advancements in comparative genomics have provided tools to expand genetic studies of convergent phenotypes based on a few candidate genes to entire genomes, and how such large-scale genomic data are being used to explore the rate and pattern of convergence at different hierarchical levels. In particular, we highlight the need to carefully define the convergent phenotype and utilizing the role of natural history records in aiding this definition. As generating genomic data becomes easier with time, integration of community-wide organismal expertise and natural history collections will remain key to understanding the genomics of convergence. We focus primarily on convergence among distantly related species, largely in animals, and generally do not discuss convergence among close relatives or populations.

### Role of organismal expertise in understanding convergence

The resurgence of interest in phenotypic convergence is driven by the desire to add a new layer – genomics - to what has been a long-standing, centuries-old interest in natural history and organismal biology (Fig. 1). Without genomics, studies of phenotypic convergence would no doubt continue as they have for decades, particularly given the firm foundation of comparative biology on which studies of convergence now rest [15,16]. However, the rapidly declining costs of genome sequencing have reinvigorated questions about the degree to which convergent phenotypes share a genetic basis, and generated considerable excitement about using convergence as a means to understand the genetic basis of phenotypes [5]. However, the “genomics gold rush” in studies of convergence has tended to focus on a few easily defined and extensively studied traits, such as the transition to marine [17], subterranean [18], or high-altitude [13] life, loss of flight in birds [19], eusociality in insects [20], social behavior in vertebrates [21,22], vocal learning in birds [23], echolocation in mammals [14,24,25], among others. While these studies have laid the groundwork for the field, organismal and natural history expertise remains critical for the maturation of studies relating phenotypic convergence and genomics.

#### Definition and complex nature of convergent phenotypes

Organismal expertise and knowledge of natural history data can inform comparative genomics in several ways. A mechanistic understanding of convergent phenotypes ultimately requires in-depth knowledge of how organisms function in the wild. It is relatively easy to designate a given species as having either a subterranean lifestyle or a lifestyle wholly above ground, but such a simple dichotomy might mask the substantial diversity of ecological and behavioral traits even within the subterranean lifestyle – for example, the diversity of burrow structures and whether or how the work of digging the burrow is shared between the sexes. Similarly, categorization of species as either ‘marine’ or ‘non-marine’, or ‘volant’ and ‘flightless’ will no doubt capture important components of phenotypic convergence, without necessarily advancing a mechanistic understanding of these phenotypes. Such categorization ignores behavioral, developmental, physiological and ecological complexity that will add nuance to any comparative analysis. For example, birds categorized as “flightless” may exhibit forelimb morphologies varying from complete absence (e.g., moas, *Hesperornis*) to slightly shortened (e.g., ostriches and Galapagos Cormorants) to highly modified forelimbs actively deployed in diving underwater but not in flight (e.g., penguins, Great Auk; Fig. 2a). Similarly, “limblessness” in squamate reptiles can mean complete loss of forelimbs, hindlimbs or both, or partial loss of digits and/or limb long bones (Fig. 2b). While simple binary categorizations of specific character states have proven powerful in guiding comparative genomic analyses [18], finer dissection of convergent phenotypes as a quantitative continuum rather than a binary phenomenon [26] will allow both an adequate testing of the adaptive value of the traits in question and a more detailed categorization of adaptations themselves. Recent models designed to test the significance of genotype-phenotype associations in a phylogenetic context are an important part of this new framework [27,28]. An important question is how to derive the most statistical power to detect convergent genotype-phenotype associations when phenotypes are defined continuously or with greater than two states. New quantitative methods of assessing convergence in phenotypic traits [29–31], as well as phylogenetic quantitative genetic models [32,33] will both be helpful in accommodating complex characters into genomic studies of convergent evolution.

**Fig. 2.**
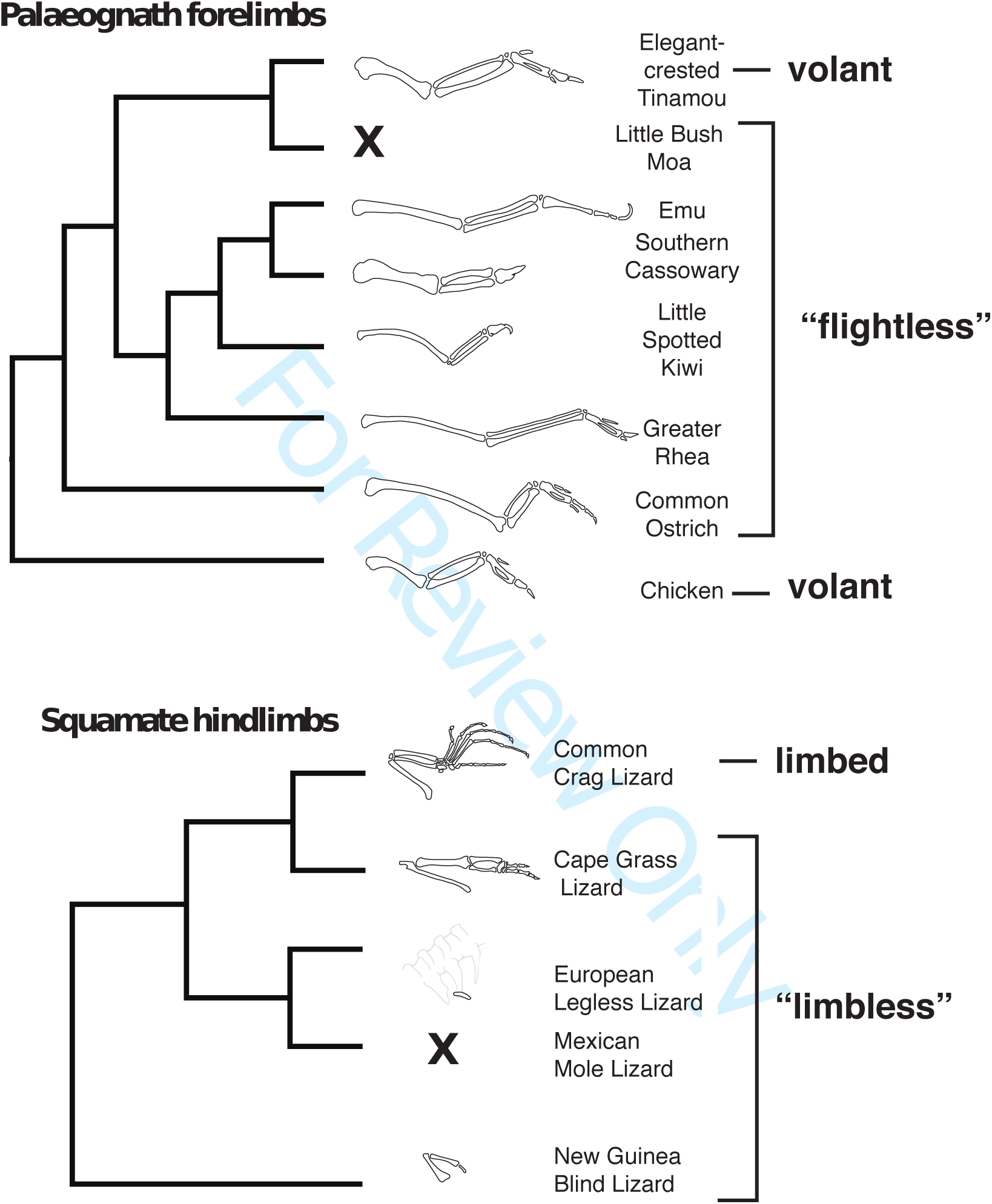
Illustration of the continuous nature of limb states often categorized as binary in studies of evolutionary convergence. In both panels, taxa were chosen to illustrate a range of forelimb character states and for which we had easily verified details from microscopy, photographs or specimens. X indicates complete absence of fore-or hindlimb elements. Drawings not to scale. Top, tree for palaeognathous birds (topology after [18]), with representative drawings of forelimbs for taxa casually deemed volant or flightless. Taxa and sources as follows: Elegant Crested Tinamou (*Eudromia elegans*, MCZ343064 and 340325); Little Bush Moa (*Anomalopteryx didiformis*; all forelimb elements absent); Emu (*Dromaius novaehollandiae*; after photo by JAC from Muséum National d’Histoire Naturelle [MHNH]); Southern Cassowary (Casuarius casuarius, JAC MHNH photo, MCZ364589); Little Spotted Kiwi (*Apteryx owenii*, MCZ340308); Greater Rhea (*Rhea americana*, JAC MHNH photo, MCZ341488); Common Ostrich (*Struthio camelus*, JAC MHN photo, MCZ341420); Chicken (*Gallus gallus*, online sources). Bottom, examples of limbed and limbless squamates. Relationships after [104]. Common Crag Lizard (*Pseudocordylus melanotus*, CAS173019); Cape Grass Lizard (*Chamaesaura anguina*, MCZ R-173157); European Legless Lizard (*Pseudopus apodus*, CAS-184449), body cavity shown in light gray to provide positional context for hindlimb; Mexican Mole Lizard (*Bipes biporus*, CAS-142262, hindlimb absent); New Guinea Blind Lizard (*Dibamus novaeguineae*, CAS-SU 27070). MCZ, Museum of Comparative Zoology, Harvard; CAS, California Academy of Sciences. All limb drawings by Lily Lu.

#### Diverse types of natural history knowledge can inform comparative genomic studies of convergence

Whereas the above perspective emphasizes ‘vertical’ knowledge, that is, deep understanding of the natural history of individual species, comprehending the breadth of taxonomic diversity across clades is a second way in which natural history knowledge can inform comparative genomics. Integration of this ‘horizontal’ knowledge of the total biology of a particular clade of organisms will be important to broaden our perspective on convergent evolution (Fig. 1). The wealth of convergent traits across the Tree of Life is likely to be found not in textbooks but in taxonomic monographs written by naturalists and curators over the last couple of centuries [34]. One example of such a convergent trait is testis color in birds: why are testes in disparate groups of birds black instead of the usual tan? Another example is the evolution of parity mode (oviparous or viviparous), which otherwise being highly conserved trait in amniotes, shows complex mosaic of convergence in squamates. The number of such convergent traits is seemingly limitless, yet we know little about the power of comparative genomics to unravel the molecular basis of these phenotypes, or how our understanding of the link between genotypic and phenotypic convergence will change as the number and type of convergent traits studied from a genomic perspective increase.

Natural history and phenomic knowledge from fossils can inform interpretation of character polarity, diversity and variation by directly informing the number and type of occurrences of convergence in extinct and extant taxa. For example, the number of extant flightless avian taxa is much smaller than the number of flightless avian taxa known from the fossil record, with many convergent instances of flightlessness represented only by extinct taxa (e.g. elephant birds, moa, adzebills, the Atitlán Grebe, the Great Auk, the Kaua’i Mole Duck [35]. The inclusion of fossil taxa near the base of clades can clarify whether traits are derived and potentially convergent or with a single origin and ancestral to a group. This may provide a better estimate of the ancestral phenotype from which the convergent traits evolved. Fossil data from natural history collections are similarly crucial for calibrating phylogenies by time, allowing investigations to assess not only which phenotypes are convergent, but when they arose. In addition, divergence time data among various taxa available in public database (e.g. TimeTree [36]) provide useful information for reconstructing ancestral. The temporal data provided by fossils are important to assessing the viability of potential causal hypotheses of drivers of convergent phenotypes. Recently extinct taxa may also provide genomic data, allowing direct incorporation into molecular phylogenies with extant taxa [37,38].

Museum collections, and the wealth of phenotypic data that they provide, are an excellent source of natural history knowledge [39,40]. Museum specimens are critical for verifying species identities, claims of specific phenotypes published in the scientific literature, are the primary source for scoring characters not yet explored for many species [41], and document diverse aspects of organismal phenotypes including anatomy, environmental context, and various types of nanostructures and chemical profiles. For instance, thorough phenotypic revisions of museum specimens have confirmed the existence of taxonomic misidentifications leading to the description of new species of Cotinga *(Tijuca condita)*, previously misidentified as *Tijuca atra* [42], or a more comprehensive understanding of phenotypic diversity and conservation needs of endemic Neotropical procyonid genus *Bassaricyon* [43]. Natural history records also provide vital information for interpreting downstream results from genomic analysis. Therefore, it is crucial that published genomes should when possible be based on DNA derived from a traceable, documented source, such as a vouchered museum specimen, known lab variant or strain, captive animal, or lab colony. In cases where this is not possible (e.g., a wild-caught individual that is entirely destroyed in the process of extracting DNA), imaging and documentation of provenance as well as associating as much additional metadata as possible is still crucial.

Many good examples of integrative comparative genomics investigations of convergence come from research teams that includes curators, taxonomists, naturalists or other experts in organismal diversity in morphology, function, and ethology at universities or other research settings [44]. However, a cursory analysis of keywords and author addresses in the Web of Science suggests a paradox: whereas museum scientists have frequently published on the general topic of evolutionary convergence, they now appear underrepresented in the second wave of convergence studies based on comparative genomics (Fig. 3). We recognize that relevant expertise in organismal phenotypes is also housed in great abundance in diverse university settings without affiliated museum collections, and our simple analysis will not capture and likely grossly underestimates these contributions. Nonetheless, we predict that as genomic studies of convergence mature, museum scientists, with their expertise in taxonomy, morphology, ecology, and biogeography will play an increasing role in studies of the genomics of convergence.

**Fig. 3.**
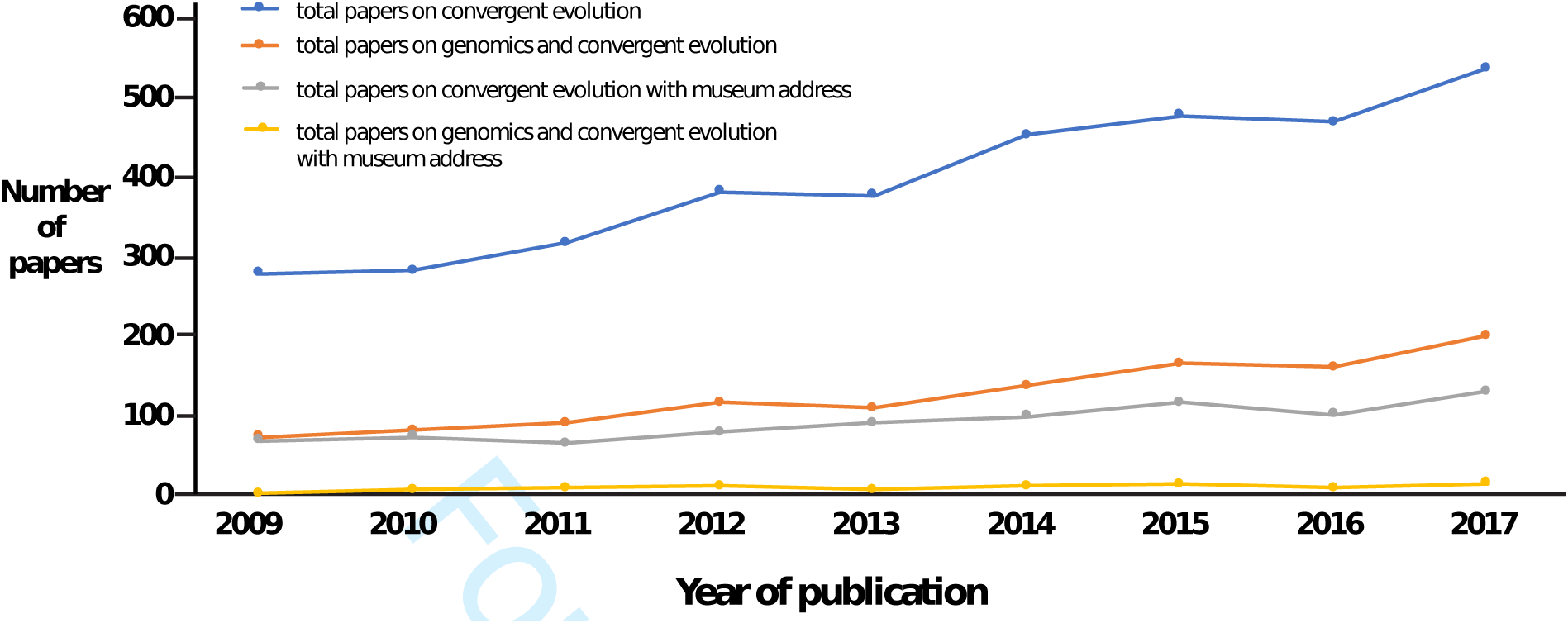
Numbers of papers on different kinds of evolutionary convergence by authors with or without a museum address (Search 1, see below). The goal of the searches was to determine if scientists with extensive natural history knowledge of organisms were participating in the second wave of studies on convergence informed by genomics (see text). We reasoned that museum specialists would constitute an important component of this community of researchers. We therefore conducted two searches on the Web of Science Core Collection on September 23, 2018 (Searches 1 and 2) and two searches on Pubmed (searches 3 and 4), on the same date. For each database we conducted two searches: Searches 1 and 3: Topic: “convergen*” AND “evolution” without or with, respectively, “genom*” as an additional topic keyword. To determine which papers had authors with museum addresses, we included “museum” OR “musee” or “museo” as part of the author address. For searches 2 and 4, we used “convergent evolution” OR “parallel evolution” as topic keywords, again, without or with, respectively, “genom*” as an additional topic keyword. Museum addresses were determined as in searches 1 and 3. The graph and associated data (Supplementary Table 1) suggests that researchers with a museum address publish extensively on general evolutionary convergence, appearing on between 7.0% and 24.2% of papers in this literature depending on the search terms. However, researchers with a museum address appear on only 4.9% to 7.0% of papers on genomics of convergence (Supplementary Table 1). These addresses are underrepresented on papers on genomics of convergence by 29%-71%, depending on the analysis. We recognize that our search is likely to miss many individuals with extensive knowledge of organismal diversity that do not work in museums or have a museum address on their publications. Additionally, our search terms are likely to detect many papers that are tangential to this analysis (see Suppementary Table 1). Nonetheless, we suspect that the trends indicated reflect the coarse-grained approach to phenotypes that have partly characterized the second wave of studies on evolutionary convergence informed by genomics. We predict that the actual numbers of authors with extensive natural history knowledge, irrespective of their work addresses, and who have participated in studies of the genomics of convergence, would change the slopes but not relative magnitude of the trends seen here.

Another way in which natural history knowledge can inform comparative genomics is through a relatively recent type of natural history – the natural history of genomes and physiological and biochemical pathways. We regard any deep knowledge of organismal function across diverse clades of organisms as a type of natural history knowledge. A good example of this type of knowledge is our understanding of the taxonomic distribution of the ability to synthesize vitamin C [45]. This case was used to powerfully demonstrate how comparative genomics can help pinpoint the likely genomic basis of convergent traits, in this case the convergent loss of the ability to synthesize vitamin C. Such biochemical knowledge was amassed through measurement by diverse laboratories of vitamin C levels in diverse organisms (in this case, from 20 separate publications spanning 1956 to 2003). Another example is the convergent ability of insects to feed on toxic plants, which is mediated by convergent substitutions, duplications and gene expression changes in a gene called ATPα [46]. Such detailed biochemical knowledge has been a major driver of recent studies of the genomics of convergence. Such an approach is likely to be a powerful method for understanding the genomics of convergence because the association between genotype and phenotype in such adaptations is likely to be tight and involve few genes, and in addition had a clearly defined convergent phenotype. Many studies on the genomics of color in mammals and in squamates provide additional examples of a trait whose molecular basis was aided by knowledge of biochemical pathways across diverse groups of organisms [47–49]. Some molecular and biochemical traits, like genome size [50], proteins and DNA sequences, are organized into well-curated databases, but many such traits, such as vitamin C synthesis, are scattered in the literature. Given what seems like the relative ease of finding the genomic basis for such traits through comparative genomics, it will be important in the future to assemble databases of phenotypic traits that span the gamut from organismal to biochemical and physiological knowledge e.g. [51–53]. Such databases can rapidly accelerate discovery of the genomic bases of convergent traits.

**Table 1:** Examples of studies identifying genomic signals of convergence at different hierarchical levels.

| **Convergent phenotypes studied** | **Methods used** | **Main findings** | **Level of convergence identified** | **Reference** |
| --- | --- | --- | --- | --- |
| High altitude adaptation in hummingbirds | Comparative genetics | Convergent amino acid substitutions | Amino acid | [130] |
| Pseudothumb and bamboo diet in Giant Panda | Comparative genomics | Convergent amino acid substitutions | Amino acid | [131] |
| Skin coloration in lizards | Cell-based assays | Convergent amino acid substitutions | Amino acid | [49] |
| Vocal learning in birds | Comparative genomics | Convergent accelerations in genes | Gene | [132] |
| Eusociality in bees | Comparative transcriptomics | Convergent accelerated evolution in genes | Gene expression | [20] |
| Vitamin C synthesis in mammals | Comparative genomics | Gene duplication/loss | Gene | [45] |
| Loss of flight in birds | Comparative genomics | Convergent rate shifts in noncoding DNA | Regulatory | [19] |
| Transitions from solitary to group living | Comparative genomics | Increase in potential for gene regulation and decrease in diversity and abundance of transposable elements | Regulatory | [133] |
| Electric organ in fish | Comparative transcriptomics | Convergence in similar transcription factors, developmental and cellular pathways | Regulatory | [134] |
| Eusociality in insects | Comparative transcriptomics | Convergent expression in biological pathways | Regulatory | [135] |
| Evolution of stripe patterns across cichlid fish radiations | CRISPR-Cas9 Genome editing | Regulatory changes of the gene act as molecular switches | Regulatory | [107] |

### Genomics of convergence

#### Outstanding questions in the genomic study of convergence

Apart from the major aim of identifying the genomic or molecular basis of convergent traits, a great diversity of questions also motivate the study of convergent traits. For instance, how does the frequency of convergence change across hierarchical levels and does it differ appreciably at the phenotypic and molecular levels? Does a nonrandom subset of genomic changes explain most instances of convergent evolution, priming convergent evolution to be more likely in certain circumstances? Are such genomic changes more likely to be regulatory or encoded by proteins? Does standing genetic variation or *de novo* mutation account for most examples of convergent evolution? Answers to such questions will not only provide critical information about the genomics of convergence but will also contribute greatly to our understanding of adaptation and evolution in general.

#### Building comparative genomics resources to study convergence

Once the phenotype of interest is defined, the next most important experimental step in comparative genomics is producing an adequate genomic foundation for downstream work (Fig. 1). While many questions of interest can be investigated with publicly-available data only, if new genome(s) are essential for a study, choosing the optimal methods for generating genomic resources is crucial. Access to high quality samples is often a limiting factor for many nascent projects, a difficulty that can be overcome in part by ensuring high-quality tissue collections are prioritized at natural history repositories. High-molecular-weight DNA is critical for modern genome sequencing technologies and in many cases, vouchered tissue samples stored in ethanol will not suffice. In addition, proper sampling procedures in the field are the most critical step to ensure high-quality DNA for building genomic resources. As an example, immediately flash-freezing tissues in liquid nitrogen is ideal given that critical molecular information (namely RNA) is quickly degraded at higher temperatures, but in many instances, tissues are transferred to liquid nitrogen after substantial delay or deep cryofreezing is logistically not possible. Though not glamorous, more detailed investigations of tissue preservation practices are vital for enabling genomic investigations, and we advocate that natural history museums should undertake a concerted, transparent effort to create best-practices recommendations for tissue samples that mirror practices already in place for whole-organism preservation [54]. For example, fresh blood stored unfrozen in Queen’s lysis buffer [55] at 4°C has provided higher quality DNA from nucleated avian blood cells [56] than museum-grade frozen tissues, and has improved sample collection practices in the Department of Ornithology at Harvard’s Museum of Comparative Zoology.

Genome assembly contiguity and gene annotation quality are also critically important for addressing target questions and maximizing the utility and availability of data from rare tissues from natural history collections. [57,58]. Chromosomal-level genome assemblies will allow us to understand the accurate location of genes associated with phenotypic traits across the genome and better understanding of cis- and trans-regulatory factors linked to those phenotypes, as well as ensure near-complete representation of genes in the assembly [59]. For example, a recent study of 78 bird genomes found that approximately 15% of avian genes had been overlooked during genome annotation, mostly due to the effects of GC-biased nucleotide composition [60]. By accounting for these missing genes, the researchers confirmed the expected positive relationship between rates of protein evolution and life history traits like body mass, longevity, and age of sexual maturity that had been previously missed [61,62].

#### Molecular convergence in protein evolution

Many recent studies of convergence have focused on protein or codon alignments to identify amino acid positions that have convergently changed in species that share a convergent trait [63–65]. In some circumstances, conflicting placement of convergent phenotypes between gene trees and species tree can be used to identify potential genomic convergence [66, 67]. However, phylogenetic clustering of a particular gene tree can also be a product of other evolutionary or experimental processes, and further analyses are required to confirm that parallel selection in distinct taxa have led to molecular convergence. Indeed, an early study that used phylogenetic signal to identify genes convergently evolving in echolocating mammals [14] was quickly met with sharp criticism [68,69], because such phylogenetic signal could arise stochastically from biased mutational spectra rather than natural selection. Castoe et al. [70], on the other hand, found that a phylogeny based on whole mitochondrial genomes that clustered snakes with agamid lizards, a relationship unsupported by nuclear gene sequences or morphological data, was actually due to strong, convergent protein evolution in just two mitochondrial genes, producing an overwhelming, yet incorrect, phylogenetic signal.

Another approach for investigating convergent protein evolution is examining rates of protein evolution across branches of a species tree and isolating instances where accelerated rates of evolution occur independently on branches leading to organisms with convergent phenotypes [17,18,71]. Such methods utilize amino acid distance trees that are normalized by average divergence rates across the genome for each tree branch and estimate the correlation between relative evolutionary rates of genes and the evolution of a convergent trait across a phylogeny [17, 72,73]. Such rate estimates as well as ancestral reconstructions have been used to detect classic examples of convergent protein evolution, such as substitutions in Na+, K+-ATPase enzymes of herbivorous insects that mediate resistance to toxic, plant-derived cardenolides [74,75] and substitutions in voltage-gated sodium channel proteins in reptiles, amphibians and fish that mediate resistance to tetrodotoxin [76,77]. Traditional methods of measuring rates of protein evolution, such as those employing the ratio of nonsynonymous to synonymous substitutions per site (dn/ds) [72], are also useful, but care should be taken to ensure that ds is not saturated when comparing distantly related species.

#### Gene family evolution associated with convergence

Gene duplications can provide raw material for rapid evolutionary innovation [78], hence analyzing the structure of gene families can provide deeper insights into the evolutionary processes underlying convergent traits. [79– 81]. Phylogenetic approaches are available to estimate the rate of change in gene family sizes [82,83], and correlated rate shifts in taxa with convergent phenotypes implicate a gene family in the process of convergent evolution. For example, visual opsin and oxygen-binding globin families are known to vary in composition under varying ecological constraints, and convergent patterns of opsin and globin family turnover have occurred in jawed and jawless vertebrates [84,85]. Studies of venom genes across animal kingdom, which form complex protein cocktails used for capturing prey and defense, show that similar protein families are commonly co-opted into hyper-mutable venom gene arrays [86,87].

#### Relative importance of regulatory regions in convergent evolution

Because many studies on convergent evolution have focused on protein-coding regions, the role of regulatory regions underlying convergent phenotypes is typically only well understood in few model systems, like sticklebacks [88]. It is plausible that regulatory elements are less constrained, and thus able to act as important drivers of adaptive molecular evolution by altering the timing, location, or level of expression of their target gene. Recent studies have indeed shown that changes in regulatory regions are associated with the origin of key innovations such as feathers and hair [89,90], as well as convergent evolution of traits such as flower pigmentation [91], loss of flight in ratites [19] and ocular degeneration in mammals [18].

Whereas predicting protein-coding genes in genome sequences is made easier both by examining homologous sequence patterns and a wealth of easy to generate functional data (e.g., transcriptomes), *de novo* identification of regulatory regions poses a significant challenge. In the era of comparative genomics, sequence conservation in non-coding regions has served as a useful starting point to identify at least part of the suite of noncoding regulatory regions across the genome [92,93]. Unfortunately, the functional links between regulatory regions and the genes they regulate are often unclear, especially in enhancers that can act over long genomic distances. This uncertainty hinders our understanding of the connections between genotype and phenotype and requires additional approaches discussed below.

### Functional characterization of genomic convergence

Given a convergent trait of interest, and candidate loci generated from any number of the genomic investigations described in the sections above, an additional step in understanding the underlying genetic mechanism for a convergent phenotype is functional validation of such genomic loci (Fig. 1). Studies from diverse groups of organisms have indicated that convergence at the genetic level can result from shared regulatory, metabolic, and developmental pathways, protein-coding genes with similar functions, or even identical amino-acid substitutions within the same gene (Table 1). These analyses range in complexity and cost, and include experimental embryology and tissue culture work, and the creation and testing of transgenics.

In recent years, techniques including CRISPR/Cas9 and massively parallelized reporter assays (MPRAs) have been added to the toolkits of those researchers interested in creating transgenic organisms, or testing hundreds or thousands of non-coding variants for enhancer activity [94,95]. Although we are unaware of any studies of genomic convergence utilizing MRPAs at the time of writing, the possibility of functionally assessing thousands of candidate loci in cell lines will almost certainly prove fruitful. CRISPR/Cas9 and other genome editing technologies are considered in many cases to be the gold standard for functional testing, and a recent study on cavefish (*Astyanax mexicanus*) metabolism utilized this technology to demonstrate a convergent insulin resistance phenotype with potential medical relevance across fish and humans [96]. Populations of river-dwelling Mexican tetra have repeatedly become isolated in caves, and the resulting cavefish have convergently evolved pigment, metabolic, and visual adaptations. After analyzing candidate genes within the insulin pathway, researchers uncovered a protein coding change in the insulin receptor gene of two independent populations of cavefish that both show insulin resistance and larger body size compared to their surface-dwelling relatives. This same substitution was also identified in insulin-resistant humans that suffer from Rabson–Mendelhall syndrome, and when placed into a zebrafish background using CRISPR/Cas9, this amino acid change resulted in insulin-resistant zebrafish that were larger than their wild-type siblings [96].

In contrast to this transgenic work, another example of non-model vertebrate convergence in pigeon feather crests was functionally tested with *E. coli* and selective media. Following population genetic analyses that identified a substitution in the gene *EphB2* as a likely candidate for the reversal of feathers on the head of pigeons, researchers discovered a neighboring missense mutation in a species of dove with a convergent crest. In a simple and inexpensive assay, wild-type and convergently-crested *EphB2* genes were transformed into bacteria and plated; based on the known toxicity of wild-type EphB2 protein to bacteria, it was possible to determine that both crested *EphB2* convergent mutations negatively altered the protein’s biochemical function [97].

In the past, there has been considerable debate about whether the link between genotype and phenotype is explained by only few major core genes, or whether it is due to accumulation of small-effect changes at multiple loci across the genome, highlighting the important distinction between polygenic vs Mendelian phenotypes [98–100]. Similarly, there has been growing debate about whether most of the genetic variance is hidden as numerous rare variants of large effect or common variants of small effect [101]. In addition, pleiotropy (involvement of the same genes in multiple traits) poses challenges in associating a particular genetic locus with a phenotypic change [102]. Existence of pleiotropy in complex traits has been widely reported in genome-wide association studies (GWAS) [103], and this observation has been a constant challenge for evolutionary-development (evo-devo) studies [104]. Patterns of pleiotropic variants may confound linking genotypic signatures to a particular trait and systematic approaches are required to identify pleiotropic variants and their associations to infer molecular mechanisms shared by multiple traits [105]. For example, pigmentation has been widely used natural trait to assess the importance of convergent evolution at genetic level with *Agouti* and *MC1R* being identified as obvious candidate genes to have strong effect on pigmentation in vertebrates [106]. But these convergence in pigmentation have been identified at multiple levels of mutations, gene or gene functions [48,107], an example that highlights the underlying challenge of identifying causal genetic architecture associated with phenotypic convergence.

Additionally, questions about the relative roles of regulatory vs. structural protein coding variation as the main drivers of morphological evolution are not new [108,109]. In the past, studies of only few genetic loci did not provide enough resolution to indicate preference for regulatory or protein coding changes for adaptation, but the rise in large scale genomic studies on adaptive evolution in the future will continue to address this debate. Quantitative measures of the contribution of protein-coding versus regulatory to convergent traits are also needed. Even if we had the complete catalogue of mutations underlying a convergent trait, how would we quantify the relative contributions of these two mutational sources to the convergent phenotypes? Phylogenetic analogues to QTL mapping, which could provide estimates of the proportion of trait similarity between species that can be attributed to a given locus, are perhaps a distant goal, but new perspectives on quantitative genetics in a phylogenetic context are already providing glimpses of this future [33,110].

The future of convergent genomic analyses will make use of these complementary functional genomics and analytical techniques for improved resolution of the genetic architecture underlying trait evolution. Thus far, genomic analyses have affirmed that convergence does exist at the phenotypic and molecular levels, with evidence of both protein-coding and regulatory convergence down to the level of single nucleotide mutations (Table 1). A more pressing question is to what extent can functional confirmation of the effect of a given mutation close the explanatory gap between historical scenarios and molecular mechanisms? Does demonstration of a functional effect of a mutation mean that the historical sequence of mutational events has been confirmed? Depending on the experimental and historical context, functional testing of a given mutation today may or may not confirm a specific sequence of mutational events in the past.

### Future directions

Renewed interest in the study of evolutionary convergence abounds and is driven in part by the emergence of genomics. As anticipated, genomic data have yielded several examples of convergent genotypic or molecular evolution, many of which are cited in the sections above. However, excitement arising from this area of research has unfortunately overshadowed the reality that most trait classifications are quite broadly defined, resulting in incomplete or potentially biased interpretations of results. Studies of convergence will benefit from having multiple replicates of independent convergences [19] and clear hypotheses and definitions of the phenotypic traits undergoing convergence [66]. It remains challenging to identify instances of convergence for which a genomic perspective will likely lead to significant new insights.

A detailed and nuanced interpretation of phenotypic diversity will be greatly facilitated through the continued support of natural history investigations and extensive and comprehensive digitization and databasing of phenotypic traits. Although the natural history literature serves as an important resource, studies on phenotypic convergence will benefit even more from direct research on natural history collections, particularly given the potential of collections worldwide to house an increasing diversity of specimen types. Emerging databases that catalog the relationships and natural history characteristics of organisms, including the Global Biodiversity Information Facility (GBIF), [111]) and the Encyclopedia of Life [52], are a promising start towards cataloging instances of phenotypic convergence. Moreover, data-rich technologies are emerging that are capable of quickly generating detailed information on natural history characteristics, such as gross organism morphology (e.g., computerized tomography; [112,113]) and environmental preferences (e.g., geographic information systems [114] and thermal imaging [115]). A concerted initiative, such as a broadening of platforms like Phenoscape [53,116], is needed in order to integrate these data with the skills and knowledge of organismal biologists, physiologists, molecular biologists, geneticists, and other stakeholders to produce detailed, hierarchical, logically coherent and searchable descriptions of organismal phenotypes. Recent efforts in large scale digitization of scientific texts, assembling phylogenomic data matrices [117] and development of automated text mining and natural language processing approaches can also facilitate high-throughput generation of phenomic datasets [118]. In addition, development of ontological phenotypic databases that contains standard terms, definitions and synonyms that can be used to describe a phenotype is also a key in generating such phenomic resources [119]. Such integration, and the computational infrastructure to allow easy access to large data sets [120], will greatly accelerate discovery of the degree, timing, and mechanisms of convergence among taxa.

Large, data-driven and taxon-rich phylogenies with comprehensive metadata attached to each taxon are a prerequisite for scaling up of genomic studies of convergence. A principle use of phylogenies is for testing macroevolutionary models [12,121] to identify whether a trait of interest is statistically associated with other phenotypic traits or with broader ecological variables; such associations can be used to support adaptive scenarios for the evolution of a trait (e.g. [122]). Numerous analytical frameworks have recently been built to address this aim, allowing increasingly complex adaptive landscapes to be modeled and associated with both continuous and discrete phenotypic traits [121]. However, a long-recognized shortcoming of model-testing approaches in this field, and in general, is the possibility that a best-fit model may still poorly reflect empirical evolution of traits across lineages [123]. One potential solution has very recent emerged called phylogenetic natural history, a framework that advocates combining model hypothesis testing with empirically-derived knowledge to better understand macroevolutionary patterns and associations [123]. Extensions of this and other approaches will be important for the continued improvement of phylogenetic comparative methods.

Phylogenies are also important for inferring evolutionary rates for regions across the genome to identify loci putatively underlying convergent phenotypes. Such analyses are widely used for analyzing protein-coding regions even before the genomic era [124], but analogous methods designed for estimating convergent rate variation in non-coding regions, such as conserved non-exonic elements, are less well developed and therefore require additional attention [28,92,125,126]. In addition, most phylogenies are only represented as purely bifurcating and phylogenetic reticulation are often not considered [127], leaving us unable to discern ‘truly convergent’ versus ‘borrowed’ traits. Moreover, our increasing ability to amass evolutionary rate estimates for thousands of genomic loci presents additional challenges of minimizing type II errors [128]. The rapid pace at which quality genome assemblies are being produced will be an important foundation for testing all genomic compartments, both coding and noncoding for a role in convergent phenotypes.

Although new approaches for phenotyping organisms are emerging, new functional genomics approaches have yet to be integrated with comparative genomics approaches [120,129]. A major challenge for the field moving forward will therefore be combining these rich forms of species-or even tissue-or cell-specific data (such as are available for the human genome) with inferences derived from cross-species genomic comparisons to functionally evaluate genomic drivers of in convergent evolution. Integrating genomics data, cutting-edge laboratory and computational techniques, and detailed, multi-level understanding of diverse natural history data will help answer fundamental questions about the propensity for convergent evolution and the genetic and molecular underpinnings of convergent phenotypes.

## Data accessibility

This article has no additional data.

## Authors’ contributions

All authors planned the organization and themes of the paper. SL lead the writing of the paper, with substantial writing from DCC, PG, JT and SVE. All authors edited and approved the final text.

## Acknowledgements

We thank Terry Capellini and Jim Hanken for helpful discussions; Nathan Clarke and one anonymous reviewer for helpful comments on the manuscript; the curators and curatorial associates of the Department of Herpetology of the Museum of Comparative Zoology, Harvard, and of the California Academy of Science, for loaning material; and Lily Lu for the drawings in Fig. 2.

